# Modularity in the Microcircuitry of the Mouse Inferior Colliculus Yields Dual Processing Streams for Auditory and Multisensory Information

**DOI:** 10.1101/572776

**Authors:** Alexandria M. H. Lesicko, Daniel A. Llano

## Abstract

The lateral cortex of the inferior colliculus (LCIC) is parcellated into two neurochemical compartments: one that comprises periodic neurochemical modules rich in GABAergic and cholinergic terminals and an extramodular matrix rich in calretinin neurons. We recently found that projections from auditory structures (auditory cortex and central nucleus of the IC) target the extramodular matrix, while somatosensory structures (somatosensory cortex and dorsal column nuclei) target the modules. What is peculiar about this finding of segregated inputs is that previous work has found that many LCIC neurons respond to both auditory and somatosensory stimuli. To investigate how these pathways interact, here we use laser photostimulation of caged glutamate to interrogate local LCIC circuits in brain slices from mouse. We found that most cell types in the LCIC receive inputs only from their home domain, but that GABAergic neurons in the modules serve as a bridge between modules and extramodular space. Further, we found that residence in- or out-of a module strongly predicted the output connectivity of that cell. These data suggest that distinct processing streams are seen in the LCIC and that GABAergic cells in modules serve to link these streams.

## INTRODUCTION

A number of brain regions can be parcellated at the subnuclear level based on differences in neurochemistry, cytoarchitecture, or connectivity. The most heavily studied of these “modular” structures include the somatosensory barrel cortex and the patch/matrix organization of the striatum (Gerfen, 1992; Petersen, 2007). The IC, a midbrain structure that is centrally positioned within the auditory system and is thought to serve as an integration hub, also exhibits modularity (Casseday et al., 2002). Specifically, the LCIC can be subdivided into modular regions characterized by dense staining for GAD67, PV, CO, AChE, and NADPH-d, and extramodular regions that are characterized by heavy calretinin labeling (Chernock et al., 2004; Stebbings et al., 2014; Lesicko et al., 2016; Dillingham et al., 2017).

These neurochemical divisions also correlate with differences in connectivity: modular regions of the LCIC receive input primarily from extrinsic somatosensory structures, while extramodular areas receive auditory inputs from the AC and CNIC (Lesicko et al., 2016). An unresolved paradox exists in this arrangement of multimodal inputs: the apparent anatomical segregation in somatosensory and auditory inputs belies a long history of physiological studies demonstrating multisensory convergence in the LCIC. Early recordings from the cat LCIC uncovered single units that respond to both auditory and somatosensory stimuli (Aitkin et al., 1978; Aitkin et al., 1981). Other studies that have examined the effect of Sp5 or dorsal column stimulation on responses to sound in the LCIC have found that the majority of units respond bimodally (Aitkin et al., 1978; Jain and Shore, 2006).

Given the evidence for the role of the LCIC in multisensory integration, it is peculiar that the somatosensory and auditory inputs to the LCIC are spatially segregated. This lack of convergence among multisensory inputs to the LCIC suggests that a secondary mechanism of integration must be present: either information from the two senses is integrated in a lower structure that projects to the LCIC, or there is communication between modular and extramodular regions of the LCIC. In the present study, we specifically investigate the latter possibility through functional characterization of the local inputs to LCIC neurons. Additionally, we use retrograde tract tracing techniques to determine whether modular and extramodular regions of the LCIC project to distinct targets.

## METHODS

### Animals

Juvenile (postnatal day 11-21) GAD67-GFP and adult knock-in mice of both sexes were used for laser photostimulation and anatomy studies, respectively. Transgenic mice were obtained from the University of Connecticut and were bred with wild-type Swiss Webster mice to generate heterozygous progeny in which enhanced GFP is under control of the endogenous GAD67 promoter (Tamamaki et al., 2003; Ono et al., 2005). Animals were screened for phenotypic evidence of transgene expression between postnatal day 2 and 7. The screening procedure involved illuminating the dorsal surface of the scalp with blue light (the excitation range for enhanced GFP) and checking for evidence of green fluorescence in the cortex, midbrain, and cerebellum. All procedures were approved by the Institutional Animal Care and Use Committee at the University of Illinois. Animals were housed in care facilities approved by the American Association for Assessment and Accreditation of Laboratory Animal Care. Every attempt was made to minimize the number of animals used and to reduce suffering at all stages of the study.

### Slice preparation

Mice were anesthetized with a mixture of ketamine hydrochloride (100 mg/kg) and xylazine (3 mg/kg) intraperitoneally and perfused transcardially with a cold slicing solution containing: 206 mM sucrose, 10 mM MgCl_2_, 11 mM glucose, 1.25 mM NaH_2_PO_4_, 26 mM NaHCO_3_, 0.5 mM CaCl_2_, 2.5 mM KCl, and 1 mM kynurenic acid (pH = 7.4). The brain was removed and 300 μm-thick coronal tissue slices throughout the IC were obtained using a vibratome. The slices were transferred to an incubation solution (126 mM NaCl, 3 mM MgCl_2_, 10 mM glucose, 1.25 mM NaH_2_PO_4_, 26 mM NaHCO_3_, 1 mM CaCl_2_, 2.5 mM KCl; pH = 7.4) and warmed to 32 degrees C for 1 hour prior to recording.

### Electrophysiology

Tissue slices containing modular regions of the LCIC were transferred to a recording chamber and submerged in an oxygenated artificial cerebrospinal fluid (ACSF) solution containing: 126 mM NaCl, 2 mM MgCl_2_, 10 mM glucose, 1.25 mM NaH_2_PO_4_, 26 mM NaHCO_3_, 2 mM CaCl, 2.5 KCl; pH = 7.4. Modular and extramodular regions of the LCIC were identified through differential expression of GAD67-GFP under blue light illumination. Cells were categorized into four groups based on whether they were found in modular or extramodular regions of the LCIC and whether they were GAD67+ or GAD67-. After identification of cell type, neurons were patched in either a single or dual whole-cell configuration 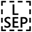 using Cs-gluconate (117 mM Cs-gluconate, 13 mM CsCl, 1 mM MgCl_2_, 0.07 mM CaCl_2_, 0.1 mM EGTA, 10 mM HEPES, 2 mM Na_2_-ATP, 0.4 mM Na-GTP) filled pipettes with tip resistances of 4-7 MΩ. Both Alexa Fluor 594 hydrazide (ThermoFisher, #A10438; 7 μM) and biocytin (Sigma-Aldrich, #B4261; 4 mM) were added to the internal solution to aid in morphological reconstruction and post-hoc confirmation of cell location. Data were acquired using a Multiclamp 700B amplifier and Digidata 1440A digitizer at a sampling rate of 20 kHz in pClamp software (Molecular Devices). Cells were held in voltage clamp at −60 mV and +10 mV to isolate excitatory and inhibitory currents, respectively, and traces were filtered with a 1 kHz Bessel filter to remove noise.

### Laser photostimulation

MNI-caged-L-glutamate (Tocris, #1490) was added to recirculating ACSF at a concentration of 150 μM and focally photolysed by a pulsed 355 nm laser (1 ms pulses). The power of the laser beam at slice level was measured and maintained at 3 mW for all experiments. The laser beam was directed into the side port of an Olympus microscope using UV-enhanced aluminum mirrors and a pair of mirror galvanometers and then focused onto the brain slice using a 10X objective. Angles of the galvanometers were computer-controlled using PrairieView software (Prairie Technologies). The Q-switch of the laser and a shutter controlled the timing of the laser pulse for stimulation. The stimulation pattern for input mapping consisted of 200 positions arranged in a 10 x 20 array, with 80 μm between adjacent rows and columns. A non-neighbor stimulation paradigm in which sequentially stimulated sites are spatially dispersed was used to prevent local accumulation of uncaged glutamate and desensitization of receptors following repeated stimulation (Shepherd et al., 2003). Excitatory and inhibitory maps were repeated 2-3 times each and averaged to ensure consistent results and reduce the effect of spontaneous inputs. To distinguish between direct activation of the recorded cell and synaptic activation of presynaptic partners, the excitatory mapping was repeated in low calcium ACSF (126 mM NaCl, 4 mM MgCl_2_, 10 mM glucose, 1.25 mM NaH_2_PO_4_, 26 mM NaHCO_3_, 0.2 mM CaCl, 2.5 KCl; pH = 7.4) to block synaptic activity, and this “direct” input map was subtracted from the original excitatory map to generate a map containing only excitatory synaptic inputs (Llano and Sherman, 2009).

### Excitation profiles

Excitability mapping was performed to determine how photostimulation at specific experimental parameters (3 mW, 1 ms laser pulses) affects the spike output of each of the four cell types of interest. Cells were patched in cell-attached mode and a 10X10 grid with 20 μm between adjacent stimulation sites was centered over the soma. For each cell, the mapping was repeated four times and the total number of spikes in the average map was compared to determine if differences in excitability exist between cell types. The average number of spikes at various distances from the soma was also computed to determine the approximate width of activation for a single laser pulse.

### Tracer injection

Mice were anesthetized intraperitoneally with a mixture of ketamine hydrochloride (100 mg/kg) and xylazine (3 mg/kg) and a small hole was drilled in the skull above the structure of interest. A glass micropipette, tip diameter 20-30 μm, was filled with a 2% solution of Fluorogold (FG) dissolved in acetate buffer (pH = 3.3) and lowered into the brain. FG was injected iontophoretically using 5 μA positive current pulses (50% duty cycle) for 10-20 minutes. A 15 μA negative holding current was applied during placement and removal of the pipette to prevent unwanted leakage of the tracer.

### Tissue processing and microscopy

After recording, slices containing biocytin-filled cells were fixed overnight in a solution of 4% PFA. Slices were rinsed three times in PBS and transferred to a solution containing 0.3% Triton X-100 and Alexa Fluor 568-conjugated streptavidin (#S-11226, ThermoFisher). To visualize cell morphology, slices were wet-mounted on coverslips and imaged using a Leica SP8 laser scanning confocal microscope and LAS X control software. Mosaic Z-stacks were taken at 20X throughout the extent of the LCIC, collapsed into 2D maximum intensity projections, and tiled into a single image.

Following a 3-7 day survival period, Fluorogold-injected animals were anesthetized with a mixture of ketamine hydrochloride (100 mg/kg) and xylazine (3 mg/kg) and perfused transcardially with 4% PFA in PBS. The brain was removed and post-fixed overnight in the PFA solution. After being cryoprotected in an ascending series of sucrose solutions, the brain was embedded and cut into 40 μm thick sections on a freezing sledge microtome. Tissue sections were imaged with a Leica SP8 laser scanning confocal microscope and LAS X control software. For each IC tissue section containing retrograde label, 20X mosaic Z-stacks were taken throughout the entire depth and x-y plane of the IC. The stacks were collapsed into 2D maximum intensity projections and tiled into a single image using LAS X software. Photoshop was used to adjust the color balance and to draw masks around the edge of the tissue to cover the embedding medium. Reconstructions and cell counts were performed using Neurolucida software. The Allen Reference Atlas was used to determine the location of injection sites and the approximate rostro-caudal coordinates of IC tissue sections (Goldowitz, 2010).

### Analysis

Custom-written MATLAB scripts were used to quantify laser-driven responses. For a given cell, a trapezoidal integration function was applied to each trace to calculate the inhibitory and excitatory charge in the first 100 ms after laser onset. These values were then converted into heat maps and overlaid with images of the GAD67-GFP fluorescence to determine if presynaptic partners were located in modular or extramodular regions of the LCIC. Each stimulation site was grouped as either modular or extramodular, and off-tissue sites were removed. The charge from responses in each of these two groups was summed to yield the total inhibitory and excitatory synaptic charge arising from modular and extramodular areas. The modularity of the excitatory and inhibitory inputs for each cell was quantified using a modularity index 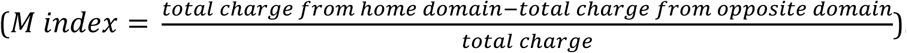. Cells with a positive index receive more input from their home domain (e.g., a cell located within a module receives more modular input than extramodular input), cells with a negative index receive more input from the opposite domain, and cells with an index near zero receive mixed input from both areas. Similarly, the balance of excitatory and inhibitory synaptic input for each cell was calculated using an E:I index 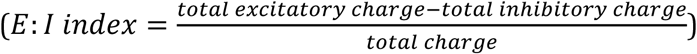.

### Statistical procedures

Shapiro-Wilk tests were used to determine if the data were normally distributed. In cases in which the assumption of normality was violated, Kruskal-Wallis and Dunn’s post-hoc testing (with a Holm adjustment for multiple comparisons) were used to compare >2 groups, and Wilcoxon rank sum tests were used for two group comparisons. For normally distributed data, one-way ANOVA and unpaired t-tests were used.

## RESULTS

### Experimental design

In order to determine the degree and directionality of communication between cells in modular and extramodular regions of the LCIC, we performed whole cell voltage-clamp recordings and stimulated pre-synaptic partners throughout the LCIC using laser photostimulation of caged glutamate. Modular and extramodular regions were visually distinguished under blue light illumination by their differential GAD67-GFP labeling in tissue slices from a transgenic mouse line (Fig. 1A). Both GAD67+ (inhibitory) and GAD67- (excitatory) cells in both regions were recorded from and voltage clamped at +10 and −60 mV to look at inhibitory and excitatory inputs, respectively (Fig. 1A, Fig. S1A). A grid of stimulation sites was centered over the LCIC and potential presynaptic partners were stimulated in a non-neighbor fashion (Fig. 1C). Responses were plotted according to the location from which they were generated and converted into heat maps by computing the area under the curve of each response (Fig. 1C, D). Excitatory mapping was repeated in low calcium ACSF, and this “direct” input map was subtracted from the original excitatory map to isolate excitatory synaptic inputs (Fig. 1E).

**Figure 1:**
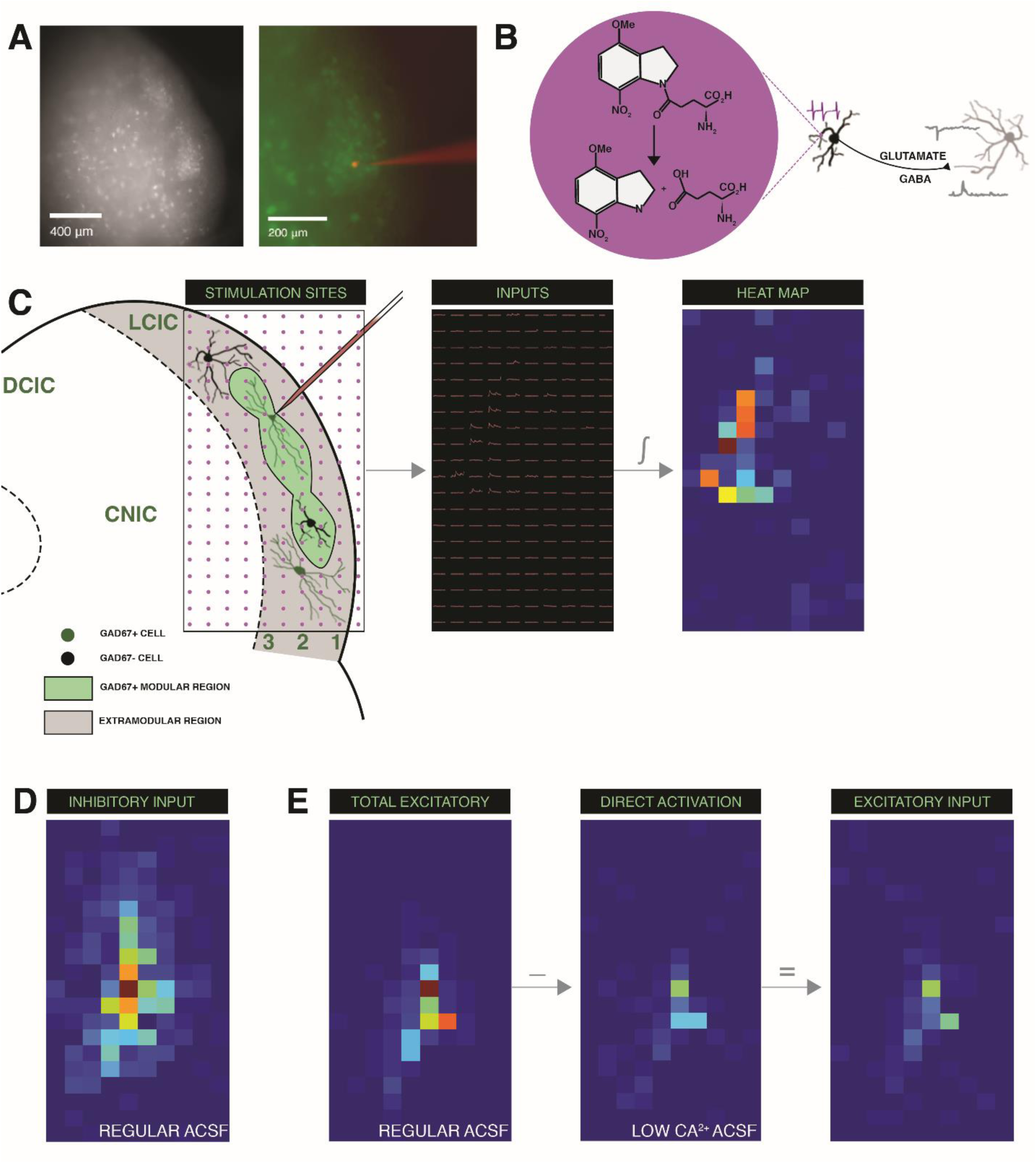
Experimental design. A) Experiments were performed in tissue slices from the GAD67-GFP mouse in which modules are visually distinguishable under blue light illumination. Cells were voltage clamped at −60 mV and +10 mV to isolate excitatory and inhibitory inputs, respectively. B) In focal regions of UV laser activation, caged glutamate is converted to active glutamate, thus generating spikes in presynaptic partners and postsynaptic currents in the recorded cell. C) Four groups of cells in the LCIC were recorded from: GAD67+ (putative inhibitory) and GAD67-(putative excitatory) cells both inside and outside of the neurochemical modules. A 10X20 grid of stimulation sites was centered over the LCIC and potential presynaptic partners were stimulated in a non-neighbor fashion. Responses were plotted according to the location from which they were generated and converted into heat maps by computing the area under the curve of each response. D) Example of an inhibitory input map. E) To generate excitatory input maps, the mapping is repeated in a low calcium ACSF to isolate direct responses. These maps are then subtracted from the map of total excitatory inputs to give the excitatory synaptic input alone.

### Excitation profiles

The spike output of cells in response to a series of laser stimuli (1 ms pulses, 3 mW) at various distances from the cell body was characterized in order to a) determine the ideal spacing between stimulation sites for experimental conditions and b) determine if the various cell types of interest differ in how strongly they are driven by photostimulation. To generate these excitation profiles, cells were patched in a cell-attached configuration and laser stimuli were delivered in a 10X10 grid (20 μm between stimulation sites) centered over the soma (Fig. S1B). The number of spikes generated in the first 100 ms after laser onset from each stimulation site was computed and converted into a heat map (Fig. S1B). Mapping was repeated four times for each cell, and the total number of spikes in the average map was summed and compared across cell types (Fig. S1C). No statistically significant differences were found across cell types (Kruskal-Wallis test, p = 0.88), though the GAD+ cells outside the modules tended to have lower spike output than the other cell types (Fig. S1C). The average number of spikes at 20 μm increments from the cell body was also determined for each cell type. Though minor differences existed in the spike output between cell types at <40 μm from the cell body, the spike output for all cells plateaued at distances further than 40 μm and dropped off to <1 spike/stimulation site (Fig. S1D). These data suggest that, at a spacing of 80 μm between stimulation sites (40 μm radius of activation), a given laser stimulus is likely driving cells in largely distinct, non-overlapping regions of the tissue. This spacing was therefore used for all mapping experiments throughout the rest of the study.

### Responses to photostimulation

For all recorded cells, input mapping was repeated for a total of three runs and averaged in order to decrease the influence of spontaneous inputs and confirm consistency across trials (note that spontaneous inputs in “Run 1” and “Run 2” of Fig. S2C are not present in the “Average” map). Large-amplitude inhibitory inputs were commonly found for all cell types (Fig. S2A). Excitatory input mapping reflected a combination of synaptic excitation and direct activation of the recorded cell (Fig. S2B). To isolate direct responses, the mapping was repeated in a low-calcium ACSF (Fig. S2C). These maps were subtracted from the total excitation maps to generate maps that reflected synaptic excitation alone (Fig. S2D). Excitatory inputs were generally smaller in amplitude and sparser than inhibitory inputs (Fig. Fig. S2A,D).

### Response patterns for cells in extramodular regions

Inhibitory input maps were generated for a total of 25 GAD- and 29 GAD+ cells in extramodular regions, and excitatory input maps were also obtained for 10 and 13 of these same cells, respectively. Heat maps were overlaid with an image of the GAD67-GFP labeling in the slice and a region of interest was drawn around the border of any modules present in the tissue. For GAD-cells outside the modules, inhibitory inputs predominately arose from stimulation sites outside of the modules, while few or no inputs were seen coming from within the modules (Fig. 2A-C). Most of the excitatory input to these cells arose from direct activation of the recorded cell, and only very sparse synaptic excitation was observed (Fig. 2A). A similar pattern of inhibitory input was seen for GAD+ cells outside the modules: virtually all of the inhibitory input arose from extramodular regions of the LCIC (Fig. 2D-F). Excitatory input maps for these cells were more heterogeneous, with some showing very sparse excitatory input (Fig. 2D), and others showing synaptic excitation that matched or exceeded the amount of synaptic inhibition (group data in Fig. 4B).

**Figure 2:**
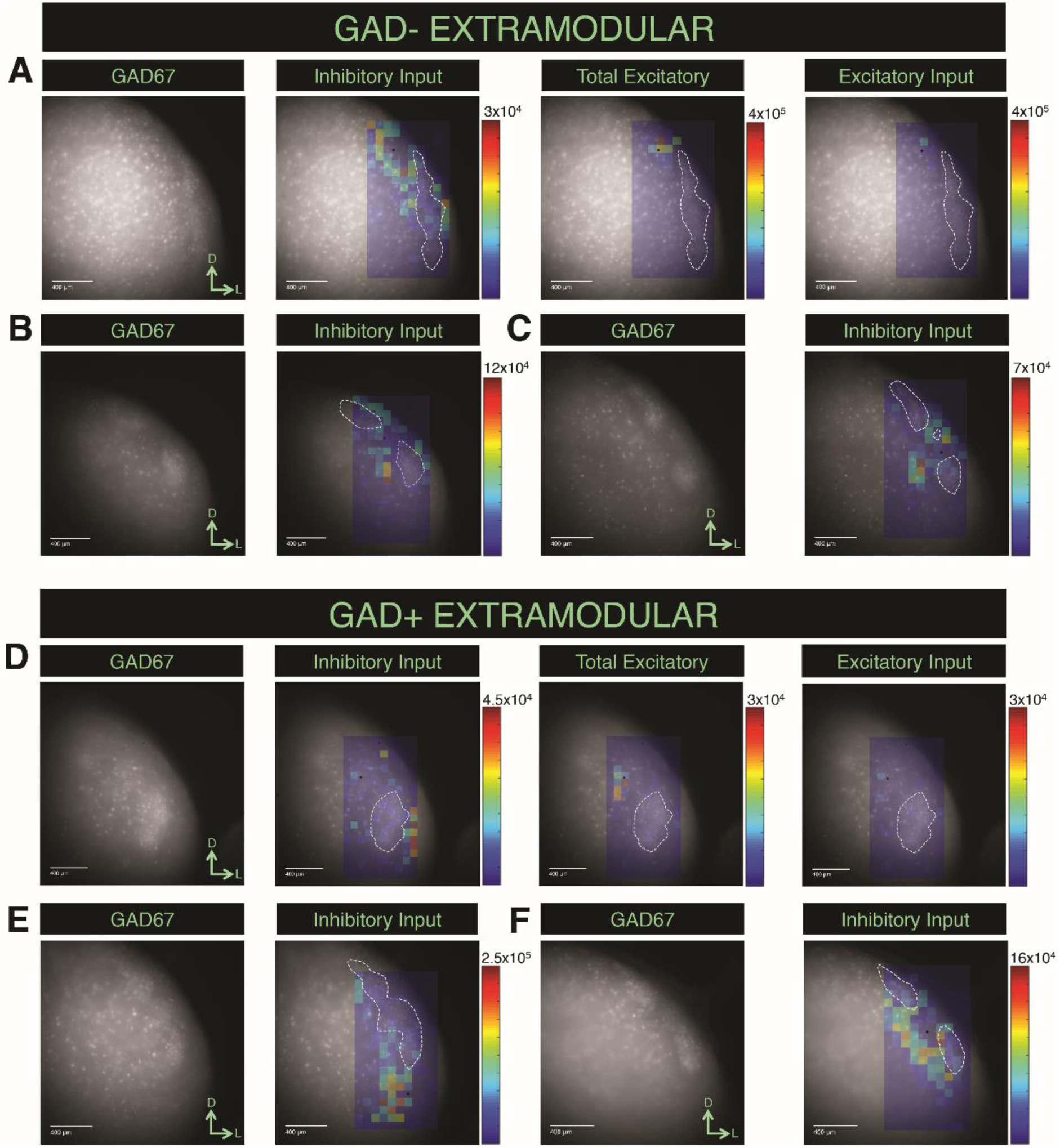
Input patterns for cells in extramodular regions of the LCIC. A) Inhibitory and excitatory data from a p12 mouse. Black dot indicates location of cell body. White outlines represent the borders of the modules. B) Inhibitory data from a p15 mouse. C) Inhibitory data from a p14 mouse. D) Inhibitory and excitatory data from a p14 mouse. Black dot indicates location of cell body. White outlines represent the borders of the modules. E) Inhibitory data from a p15 mouse. F) Inhibitory data from a p15 mouse.

**Figure 3:**
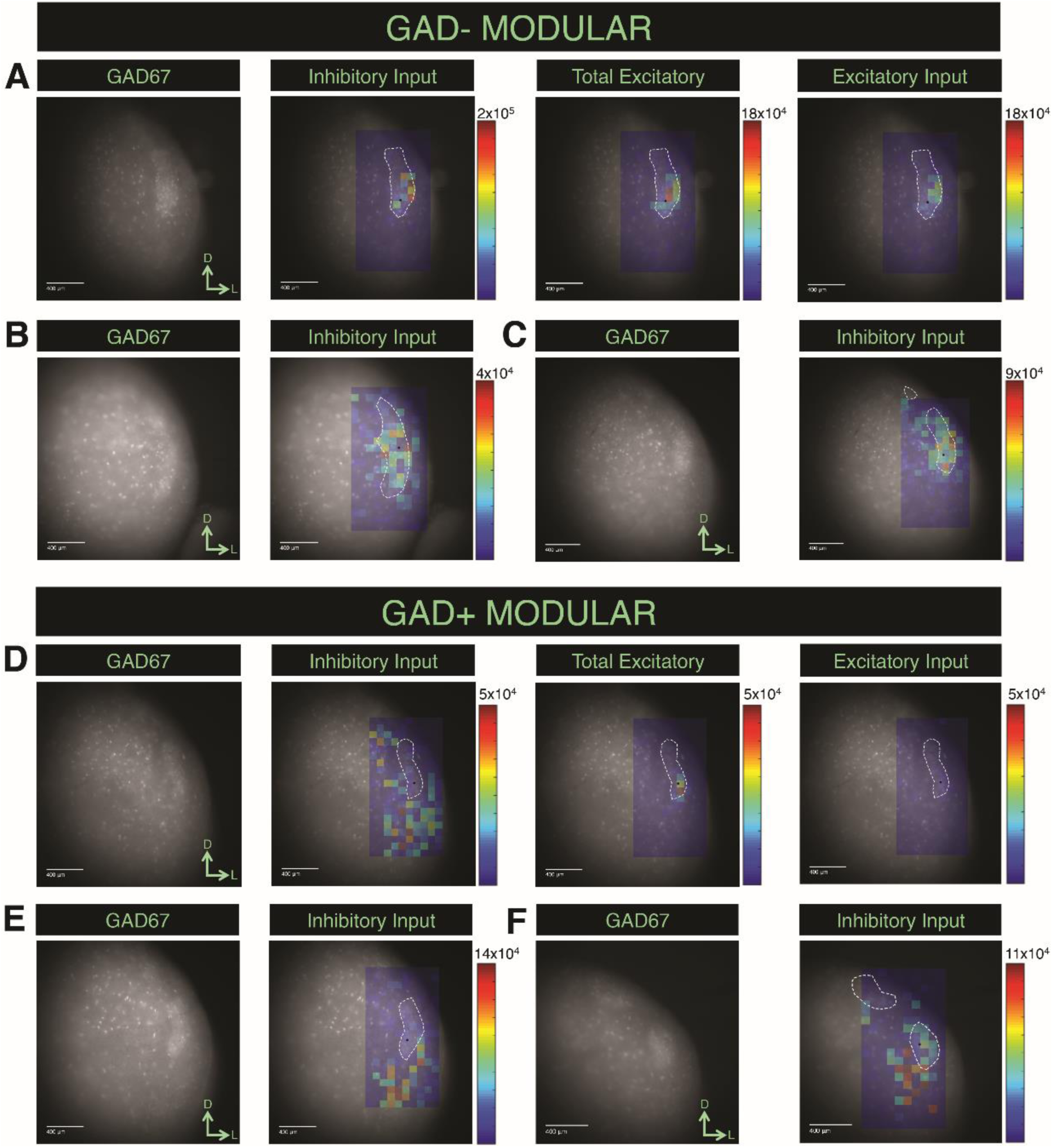
Input patterns for cells in modular regions of the LCIC. A) Inhibitory and excitatory data from a p11 mouse. Black dot indicates location of cell body. White outlines represent the borders of the modules. B) Inhibitory data from a p14 mouse. C) Inhibitory data from a p11 mouse. D) Inhibitory and excitatory data from a p15 mouse. Black dot indicates location of cell body. White outlines represent the borders of the modules. E) Inhibitory data from a p15 mouse. F) Inhibitory data from a p18 mouse.

**Figure 4:**
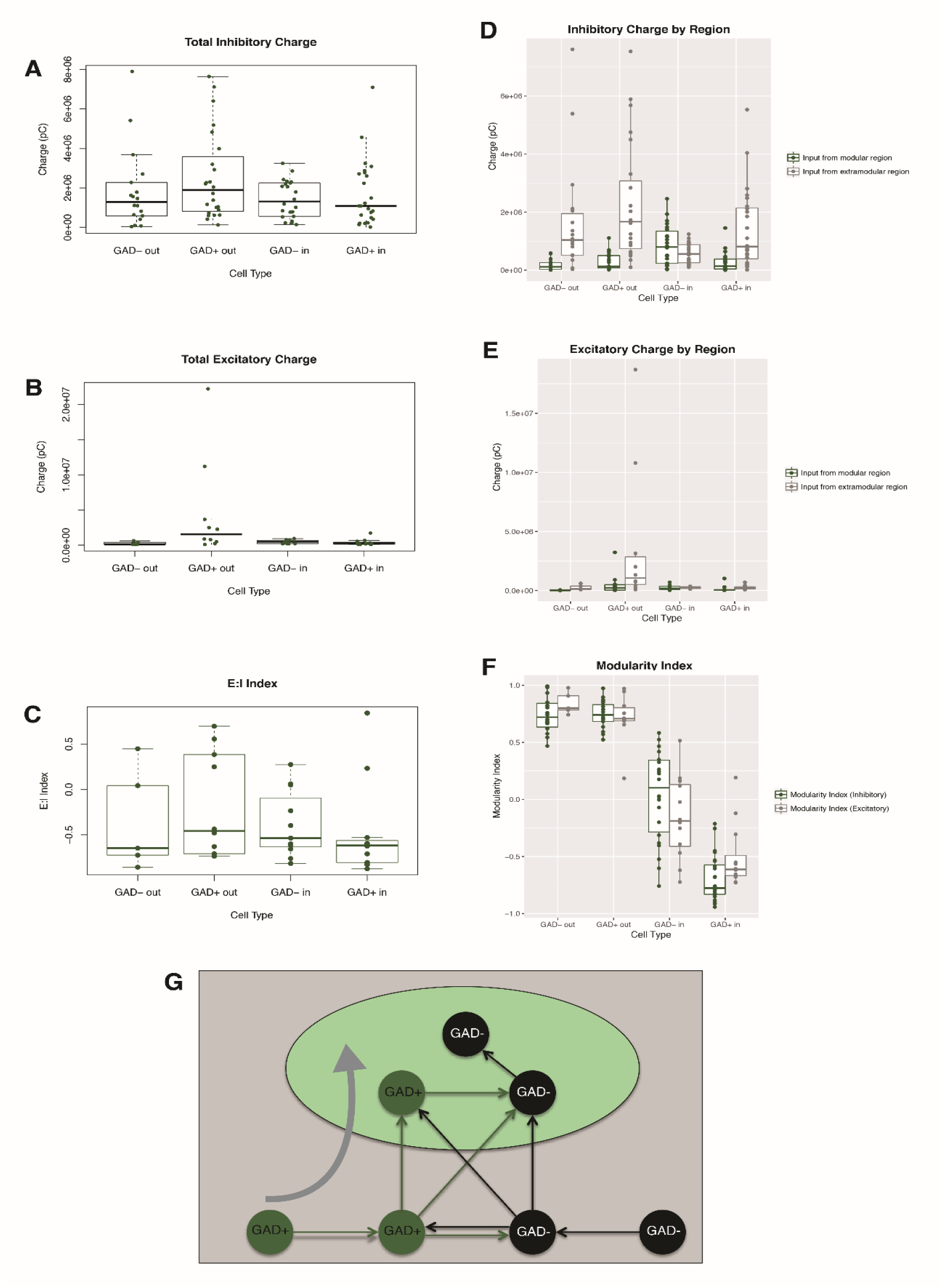
Input patterns vary according to cell type and location. A) The total inhibitory charge for each of the four cell types. B) The total excitatory charge for each of the four cell types. C) The balance of excitation and inhibition to each cell was calculated as an E:I index, and the distribution of these values for each cell type are shown as boxplots. D) Inhibitory charge coming from modular (green) or extramodular (gray) regions. E) Excitatory charge coming from modular (green) or extramodular (gray) regions. F) Modularity indices for each of the four cell types of interest. G) Cell-type specific input patterns give rise to a unidirectional flow of information from extramodular to modular regions of the LCIC.

### Response patterns for cells in modular regions

Inhibitory input maps were generated for a total of 32 GAD- and 33 GAD+ cells in modular regions of the LCIC, and excitatory input maps were also obtained for 15 of these same cells for both cell types. Some GAD-cells inside the modules received predominately clustered inputs from within a module (Fig. 3A), while others received mixed input from both modular and extramodular regions (Fig. 3B,C). GAD+ cells inside the modules showed a unique pattern in terms of their inhibitory inputs: most of the input to these cells arose from sites in the “opposite domain” (i.e. extramodular regions), with very little input coming from within the modules (Fig. 3D-F). The excitatory input to this cell type was very sparse (Fig. 3A).

### Total inhibitory and excitatory charge

The total inhibitory and excitatory charge for each map was calculated by summing the responses across all stimulation sites. The total inhibitory charge did not differ across cell types (Kruskal-Wallis test, p=0.09538), but the total excitatory charge did (Kruskal-Wallis test, p= 0.006511; Fig. 4A-C). GAD+ cells outside the modules received significantly more total excitatory charge than GAD+ cells inside the modules and GAD-cells outside the modules (post-hoc Dunn’s test with Holm correction, p=0.0034 and p=0.0199, respectively; Fig. 4B). All of the cell types except GAD+ cells outside the modules received significantly more total inhibitory than total excitatory charge (Wilcoxon rank sum tests, GAD-out: p=0.017, GAD-in: p= 0.00088, GAD+ in: p=0.020). For each cell, an E:I index was computed and the average E:I index was calculated for each cell type. Each of the four cell types had a negative mean E:I index, indicating that they are dominated by inhibition (Fig. 4C).

### Input mapping under GABA_A_ receptor blockade

To confirm that inhibitory responses are mediated via GABA, we repeated input mapping for a subset of cells in the presence of 20 μM GABAzine, a GABA_A_ receptor antagonist. Inhibitory responses were completely abolished in the presence of GABAzine, indicating that intrinsic inhibitory inputs to cells in the LCIC are GABAergic (Fig. S3A). We also repeated excitatory input mapping in the presence of GABAzine to a) determine if inhibition in the LCIC was “masking” excitatory responses and b) to confirm that our stimulation parameters elicit monosynaptic responses, since disinhibition has been shown to recruit polysynaptic circuits (Sivaramakrishnan and Oliver, 2006; Chandrasekaran et al., 2013; Sturm et al., 2014). In either instance, an increase in the overall map area would be expected. However, in the presence of GABAzine, excitatory input maps remained largely unaltered or shrank slightly in terms of their area (Fig. S3B,C).

### Charge from modular and extramodular regions

To determine if differences exist in the amount of charge coming from modular and extramodular regions, stimulation sites for each map were categorized as lying either within or outside of a module and the total amount of charge coming from each of the two regions was summed. Both cell types outside of the modules and GAD+ cells inside the modules received significantly more inhibitory input from extramodular than modular regions (Wilcoxon rank sum tests, GAD-out: p=1.68e-06, GAD+ out: p=3.49e-08, GAD+ in: p=8.5e-06; Fig. 4D). The same pattern was seen for the excitatory inputs to each cell type (Wilcoxon rank sum tests, GAD-out: p=0.00032, GAD+ out: p=0.0023, GAD+ in: p= 0.0090; Fig. 4E). The “modularity” of the inhibitory and excitatory inputs to each cell was determined using the following equation: 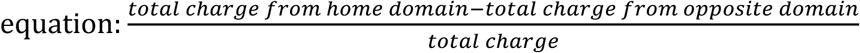. According to this “modularity index”, cells that receive the majority of their input from the region in which their cell body is located will have a positive value, cells that receive the majority of their input from the opposite region will have a negative value, and cells that receive mixed input will have a value close to zero. Both cell types inside the modules exhibited positive modularity indices for both inhibition and excitation. GAD-cells inside the modules showed evidence of mixed input, with excitatory and inhibitory modularity indices close to zero. GAD+ cells outside the modules had negative modularity indices (more pronounced for inhibition than excitation), indicating that most of their input is coming from the extramodular area (Fig. 4F).

### Paired recordings

Since clear differences were observed in the inhibitory and excitatory input patterns for cells that differed in terms of location (inside or outside of a module) and type (GAD67+ or GAD67-), we decided to further investigate how each of these factors affects the amount of common input to a given pair of cells (Fig. 5A). Paired recordings were made from a subset of 80 cells (40 pairs) that were either matched in terms of location and type (Fig. 5B bottom panel), differed in only one parameter, or were unmatched in both parameters (Fig. 5B top panel). Cross-correlations were computed between the responses at corresponding stimulation sites, and these values were compared across each group of pairs (Fig. 5C). In general, pairs that were more similar in terms of the presence of GAD or modular vs. extramodular location had larger cross-correlation values, while cells that differed in one or both parameters had smaller cross-correlation values (Fig. 5C). Cross-correlation values were significantly different between different types of pairs (Kruskal-Wallis test, p=0.016); specifically, pairs in which both the location and cell type were matched had significantly higher cross-correlation values than pairs in which both of these parameters differed (post-hoc Dunn’s test with Holm correction, p=0.0043). To determine if the distance between the cells in a pair could account for these differences, we correlated distance and cross-correlation values and found only a weak correlation (R^2^=0.0008, Fig. 5D).

**Figure 5:**
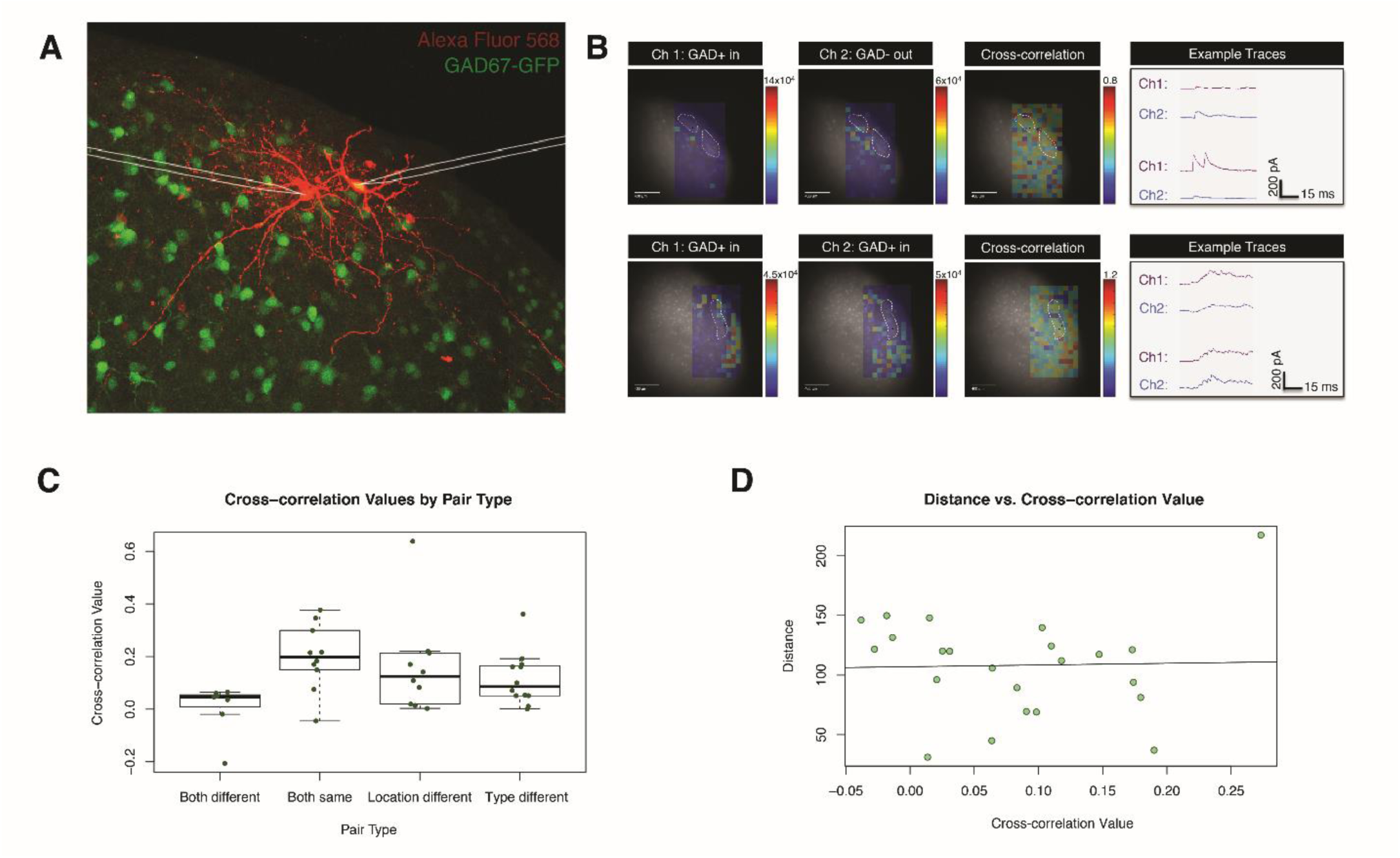
Paired recordings. A) Example of a pair of simultaneously recorded cells. B) Top panel: a pair of cells that is matched in terms of location and type. Bottom panel: a pair of cells that differs in both location and type. C) Average cross-correlation values for each category of pairs. D) Correlation between distance and cross-correlation value.

### Projections to the SC arise from extramodular regions

To determine if the distribution of LCIC cells projecting to the SC is correlated with the underlying pattern of neurochemical modularity, FG was iontophoretically injected at a mid-rostro-caudal level of the SC of the GAD67-GFP knock-in mouse. The injection site spanned much of the rostro-caudal axis of the SC and covered both the superficial and deep layers (Fig. S4A). UV illumination revealed several retrogradely labeled cells in the LCIC (Fig. 6A). The GAD67-GFP labeling in the LCIC was used to identify the location of the neurochemical modules, as well as to discern whether backlabeled cells were GABAergic or non-GABAergic (Fig. 6A). Overlay images of the FG and GAD67-GFP labeling revealed that a) cells projecting to the SC were almost exclusively found in the extramodular regions of the LCIC and b) the vast majority of retrogradely-labeled cells were non-GABAergic (Fig. 6A,B). These observations were confirmed with quantification; cell counts revealed that 97% of the retrogradely labeled cells in the LCIC were found in extramodular regions, and that 96% were non-GABAergic (Fig. 7C). Though cells projecting to the SC were found throughout the rostro-caudal extent of the LCIC, they were heavily concentrated in the rostral-most regions of the LCIC (Fig. 6B). Fewer labeled cells were found in more caudal regions of the LCIC, but the extramodular distribution of cells was conserved throughout this structure (Fig. 6B).

**Figure 6:**
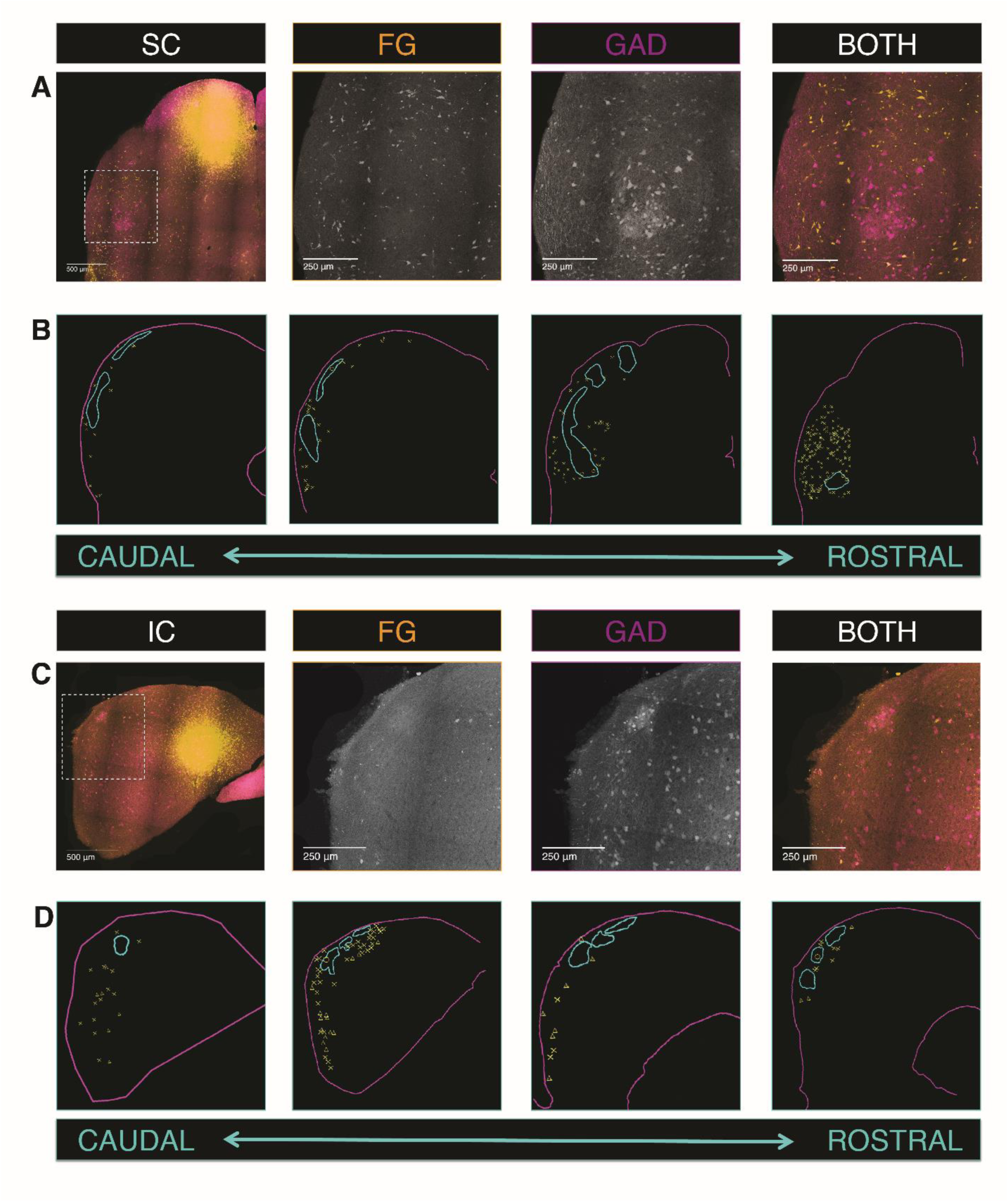
Cells that project to the SC and IC are found in extramodular regions of the LCIC. A) FG labeling in the LCIC after an SC injection in a GAD67-GFP mouse. B) Rostro-caudal distribution of cells in the LCIC that project to the SC. O: GAD-inside. +: GAD+ inside. X: GAD-outside. Δ: GAD+ outside. A) FG labeling in the LCIC after an IC injection in a GAD67-GFP mouse. B) Rostro-caudal distribution of cells in the LCIC that project to the IC.

**Figure 7:**
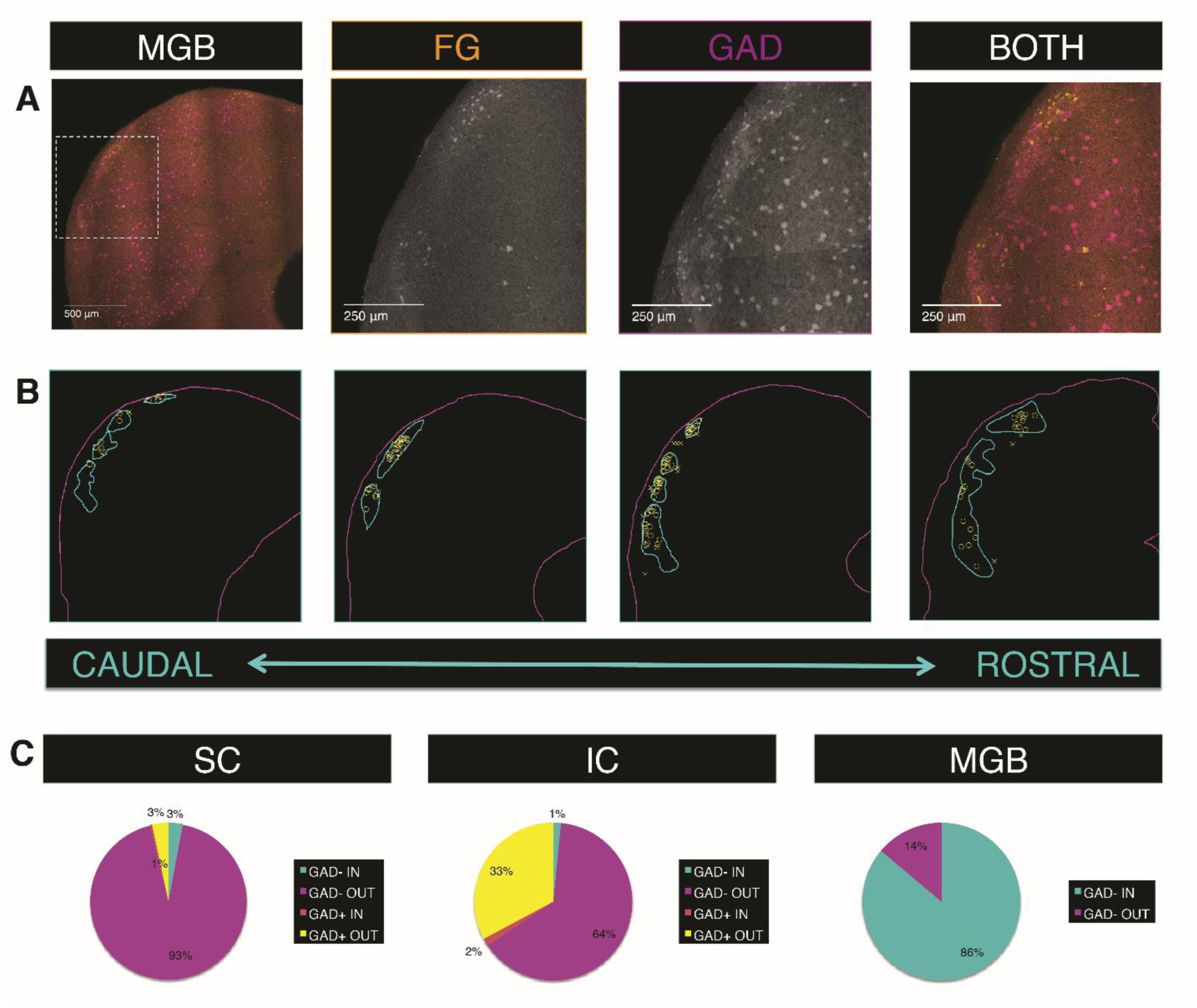
Cells that project to the mMGB are found in modular regions of the LCIC. A) FG labeling in the LCIC after an mMGB injection in a GAD67-GFP mouse. B) Rostro-caudal distribution of cells in the LCIC that project to the mMGB. C) Percentage of GABAergic and non-GABAergic cells from each LCIC subregion projecting to various targets.

### Inputs to other subnuclei of the IC come from the extramodular zone

The subdivisions of the IC are heavily interconnected, and the LCIC has been shown to project to both the DCIC and the CNIC (Coleman and Clerici, 1987). To determine if LCIC cells that project within the colliculus are also predominately distributed in either modular or extramodular regions, FG was injected into other regions of the IC. The injection site was centered in the caudal half of the IC along the border between the DCIC and the CNIC (Fig. S4B). The densest proportion of backlabeled cells was found in regions of the LCIC along the same rostro-caudal axis as the injection site (Fig. 6C,D). Overlay images showed that, similar to SC-projecting cells, cells that project intracollicularly are found almost exclusively (97%) in extramodular regions of the LCIC (Fig. 6C,D). However, a substantial percentage (33%) of backlabeled cells were found to be GFP-positive, indicating that this pathway is partially GABAergic (Fig. 6D, 7C, triangles and plus signs denote GABAergic cells).

### Cells projecting to the mMGB are found in modular regions of the LCIC

All subdivisions of the IC project heavily to the auditory thalamus, and the main thalamic target of the LCIC is the mMGB (Calford and Aitkin, 1983). A small deposit of FG was made in this region in order to backlabel colliculo-thalamic cells and determine whether their distribution in the LCIC is patterned (Fig. S4C).

According to the MGB subdivision boundaries shown in the Allen Reference Atlas, the injection site appeared largely restricted to the mMGB, with potential spillover into the surrounding paralaminar nuclei (i.e. the suprageniculate nucleus and the posterior intralaminar nucleus) (Goldowitz, 2010). In contrast to cells projecting to the SC and IC, cells projecting to the mMGB were found to form clusters that largely correlated with the location of the neurochemical modules (Fig. 7A,B). Backlabeled cells were found throughout the rostro-caudal extent of the LCIC, and their density was fairly even along this axis (Fig. 7B). Cell counts revealed that 86% of these cells were found in modular regions of the LCIC, while the remaining cells were found in the extramodular zone. Interestingly, of the 404 FG-labeled cells that were identified in the LCIC, virtually none of them were found to be double-labeled with GFP, indicating that this pathway is strictly non-GABAergic (Fig. 7C).

## DISCUSSION

In the present study, we used a combination of whole-cell patch clamp recordings, laser photostimulation of caged glutamate, and retrograde tract-tracing to determine whether modularity exists in the local connections and the outputs of the LCIC. We found that input patterns differ across cell types, with cells outside the modules receive strong extramodular input, and cells inside the modules receiving mixed or strongly extramodular input (Fig. 2,3). Cells that project to the SC were found in extramodular regions, and were found to be predominately non-GABAergic (Fig. 6). Projections to other regions of the IC also arise from extramodular areas of the LCIC, but approximately 1/3 of these cells were found to co-label for GFP, indicating that they are GABAergic (Fig. 6). In contrast to these midbrain pathways, projections to the mMGB were found to form clusters of non-GABAergic cells that largely overlap with the location of the neurochemical modules (Fig. 7).

### Cell-type specific input patterns give rise to directionality in information flow

Taken together, these input patterns suggest that cells outside the modules receive extramodular input, but send mixed output to cells both inside and outside of the modules. Cells inside the modules receive this mixed input, but relay it exclusively to other modular cells (Fig. 4G). This circuitry gives rise to a unidirectional flow of information from extramodular to modular regions of the LCIC. One advantage of such an arrangement could be to allow for a “purely auditory” computation region (the extramodular zone) that is spatially near but connectionally distinct from a multisensory computation region (the modular zone).

### Outputs of the LCIC are associated with distinct extrinsic inputs

The modular and extramodular regions of the LCIC that have been shown here to give rise to segregated outputs are also targeted by distinct inputs. Modular regions of the LCIC receive input from somatosensory structures, such as the DCoN and the primary SScx (Lesicko et al., 2016). Extramodular areas of the LCIC, on the other hand, are targeted by auditory structures such as the AC and the CNIC (Lesicko et al., 2016). Though these two subregions of the LCIC are segregated on the basis of their neurochemistry and connectivity, they do not form wholly separate processing streams; modular regions of the LCIC receive input from cells in the extramodular zone. These connections could route auditory information into the somatosensory-recipient modules, thereby forming multisensory processing zones. Extramodular regions, however, do not receive input from modular regions of the LCIC, and are therefore likely to perform computations related to auditory processing. Taken together, these data suggest that auditory information is processed in extramodular regions of the LCIC and routed to midbrain targets including the SC and other regions of the IC, while somatosensory/multisensory computations take place in modular regions of the LCIC that project to the mMGB.

### Potential pitfalls: spatial resolution of mapping

To decisively conclude whether cells in modular and extramodular regions of the LCIC communicate, our laser mapping experiments must be completed at a sufficiently high resolution to determine whether stimulation sites lie inside or outside of a given module. To address this concern, we used a 10X rather than a 4X objective for our mapping experiments, which is more commonly used (Ikrar et al., 2011; Shepherd, 2012). The UV laser spot size with this objective is 10 μm, which limits the breadth of stimulation to a more focal region. Our excitation profile experiments indicate that a given laser stimulus (1 ms in duration, 3 mW) at 10X will drive spikes, on average, within a 40 μm radius of the recorded cell. For mapping experiments, stimulation sites were spaced 80 μm apart to reduce the likelihood that adjacent stimuli would activate the same regions of tissue. It is still possible that stimulation sites on the border of modular/extramodular zones may have activated cells in both regions or in the opposite region of which the site was categorized. In this case, it may be that our interpretation underestimates the modularity of the inputs to each cell type. Regardless, our input maps and subsequent analysis show clear modularity for three of the four cell types. For the GAD-cells inside the modules, the mixed inputs from the opposite domain arise from sites that are sufficiently far enough from the border of a module for these differences to be attributed to insufficient spatial resolution (Fig. 3B,C).

### Potential pitfalls: use of juvenile animals

An additional pitfall is the use of juvenile mice, which may limit extrapolation to the adult stage. Juveniles are used since it is our experience that cell visualization for patching is very difficult beyond 30 days of age. Other investigators have found that much of the intrinsic and extrinsic connectivity of the IC is present by hearing onset (Friauf and Kandler, 1990; Gabriele et al., 2000a; Gabriele et al., 2000b). Studies that have looked at the development of the intrinsic connectivity of the CNIC have found that, while these connections undergo refinement during development, they have largely stabilized by P22 (Sturm et al., 2014). We investigated whether the E:I index and modularity index were subject to changes across the age range used in the present study, and found no significant trends (data not shown), indicating that input patterns are stable within this age range.

### Balance of inhibition and excitation

Our results indicate that all of the cell types of interest, except GAD+ cells outside of the modules, are dominated by inhibition (Fig. 4C). Since significant excitatory input was observed for at least one cell type, we can confirm that our preparation allows for the sufficient detection of excitation. It is possible, given that one of the defining characteristics of the IC is its dense inhibition, that some excitatory inputs went undetected on account of being “masked” by inhibition (González - Hernández et al., 1996). To further probe this possibility, for a subset of cells we repeated excitatory input mapping in the presence of GABAzine to block inhibitory inputs. This manipulation had very little effect on the excitatory maps overall, and in some cases led to a shrinkage of input area, rather than an expansion, which would indicate the presence of masked excitation (Fig. S3C). It is possible for photostimulation to differentially activate different cell types depending on factors such as glutamate receptor density (Shepherd, 2012). Our excitation profiles, however, indicate that excitatory cells are actually being driven more strongly by photostimulation than inhibitory cells (Fig. S1C). This indicates that the sparse excitatory input observed for most cells is not a product of insufficient activation of excitatory cells.

### Similarities to patch/matrix compartments the striatum

While the exact purpose of the neurochemical and connectional modularity found in the LCIC remains unknown, studies in other structures with a similar organization can help shed light on the potential functional advantage of such an arrangement. Modularity is also present in the striatum, with inputs and outputs being segregated according to whether they are found in the acetylcholine-rich “matrix” areas or the opiate receptor-dense “patch” areas (Graybiel and Ragsdale, 1978; Gerfen, 1984; Kincaid and Wilson, 1996). Studies that have investigated whether the two domains are fully segregated have found that the dendrites of retrogradely-filled cells in both compartments are confined to the region containing their cell bodies (Gerfen, 1985). While this suggests that the two compartments may form segregated processing streams, additional experiments have shown that intrinsic somatostatin-positive neurons form a bridge between the patch and matrix regions: the cell bodies of these interneurons are found in both regions, but their axons selectively innervate the matrix compartment (Gerfen, 1985). Single somatostatin-immunoreactive cells in the patch compartment have been found to give off axons that distribute into the surrounding matrix, suggesting that these cells provide a unidirectional projection from the patch to the matrix. Interestingly, this bears resemblance to the one-way flow of information from extramodular to modular regions demonstrated in the present study.

### Implications for multisensory processing

The results of the present study suggest a mechanism by which multisensory convergence could occur within the LCIC. Though previous studies have shown that the extrinsic multimodal inputs to the LCIC are segregated, the present study demonstrates that cells in modular regions, which are targeted by extrinsic somatosensory inputs, receive input from auditory-recipient extramodular zones (Lesicko et al., 2016). Further studies will be required to determine if single cells within the modules receive convergent input from both of these sources. It is presently unknown whether the dendrites of cells within modular and extramodular zones are confined to the region containing their soma, as is the case with other modular structures such as the striatum and the pons (Gerfen, 1985; Schwarz and Thier, 1995). If not, multisensory integration could also arise from direct input to a cell whose dendrites cross the modular/extramodular boundary. The advantage of having multisensory convergence arise from a local circuit mechanism rather than direct convergence of extrinsic inputs is presently unclear.

### Potential functional significance of modular outputs

The mMGB is the only target of cells residing in modular regions that has been identified thus far. Similar to the LCIC, this division of the auditory thalamus is known to integrate multisensory inputs, and neurons in this region exhibit broad frequency tuning and large tactile receptive fields (Aitkin, 1973; Bordi and LeDoux, 1994). The mMGB is also interconnected with limbic structures, such as the amygdala, and has been shown to mediate auditory fear conditioning (LeDoux et al., 1984; LeDoux et al., 1985). Though it has traditionally been thought that the IC provides auditory input to the mMGB, it is possible that the inputs from modular regions of the LCIC actually provide multisensory information important for executing conditioned fear behaviors (Ledoux et al., 1987).

The mMGB is also reciprocally interconnected with all regions of the AC, and it has previously been hypothesized to serve as a multisensory arousal area (Rouiller et al., 1989; Winer, 1992). Inputs from the LCIC to the mMGB could therefore convey somatosensory, auditory, or multisensory cues relevant to the animal’s state of arousal. Somatosensory convergence occurs at multiple stations within the auditory system, and has generally been thought to mediate cancellation of self-generated sounds (Wu et al., 2014). The potential participation of the LCIC-mMGB-AC circuit in this process is intriguing given that a) non-GABAergic modular cells in the LCIC, such as those that project to the mMGB, receive strong modular inhibition that could be driven by extrinsic somatosensory inputs and b) inhibiting this population of projection neurons could effectively prevent activation of auditory cortical networks and conscious awareness of self-generated noise, given that the mMGB projects widely to all areas of the AC (Rouiller et al., 1989; Winer, 1992).

### Potential functional significance of extramodular outputs

Cells in extramodular regions of the LCIC project to at least two distinct targets: the SC and other regions of the IC. It is presently unknown whether these projection systems are formed by different groups of cells, or if single cells project to both targets. Projections from the LCIC to the SC have long been thought to mediate various acoustico-motor behaviors (Huffman and Henson Jr, 1990). For example, stimulation of the IC causes movement of the pinna and eyes in conjunction with activation of auditory neurons in the SC, and this pathway is thought to mediate additional orienting and escape/defense behaviors (Syka and Straschill, 1970). Connections between the IC and the SC are also thought to be critically involved in pre-pulse inhibition of the acoustic startle reflex (Koch and Schnitzler, 1997). The SC is thought to receive information about auditory prepulses from the IC and routes this information to the pedunculopontine nucleus (PPT), a brainstem structure that also provides cholinergic input to modular areas of the LCIC (Swerdlow et al., 2001; Motts and Schofield, 2009; Schofield, 2010). The PPT then routes this information to the pontine reticular nucleus, where it converges with and influences the primary startle pathway (Davis et al., 1982).

Not only do extramodular regions of the LCIC send input to the CNIC, but they also receive dense inputs from this region (Lesicko et al., 2016). It is therefore possible that some of the LCIC cells that project to the CNIC participate in feedback loops with the lemniscal auditory pathway. In addition to inputs from the CNIC, extramodular regions of the LCIC receive descending inputs from the AC (Lesicko et al., 2016). Though descending connections from the AC to the IC predominately terminate in the LCIC and DCIC, their activation has been shown to cause striking shifts in the auditory response properties of cells in the CNIC (Andersen et al., 1980; Winer et al., 1998; Gao and Suga, 2000). Given that direct descending inputs to the CNIC are sparse, it is possible that these changes are mediated through connections from the LCIC to the CNIC (Stebbings et al., 2014).

## Acknowledgments

This work was supported by F3DC015967 to AMHL and R01DC013073 to DAL.

---

**Supplemental Figure 1:**
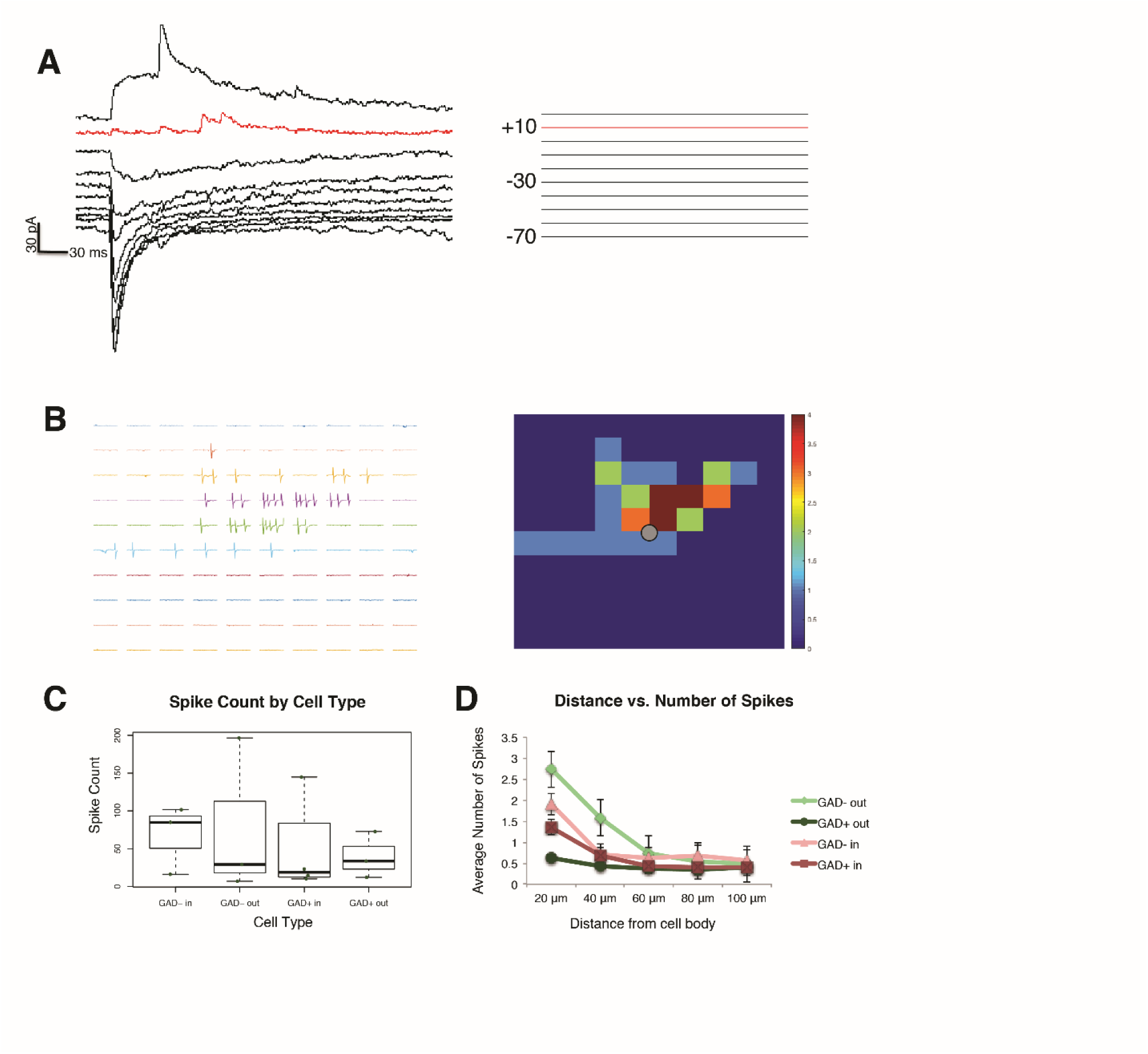
Recording and photostimulation parameters. A) Responses to photostimulation at various holding potentials. B) Left: Example of spikes recorded in cell-attached mode in response to laser photostimulation at various locations (20 μm between adjacent rows and columns) near the cell body. Right: Spike output from shown as a heat map. The location of the recorded cell is shown in black. C) The total number of spikes (average of four trials) was computed for each cell and compared across cell types. D) The average number of spikes elicited at various distances from the cell body was calculated and compared for each cell type.

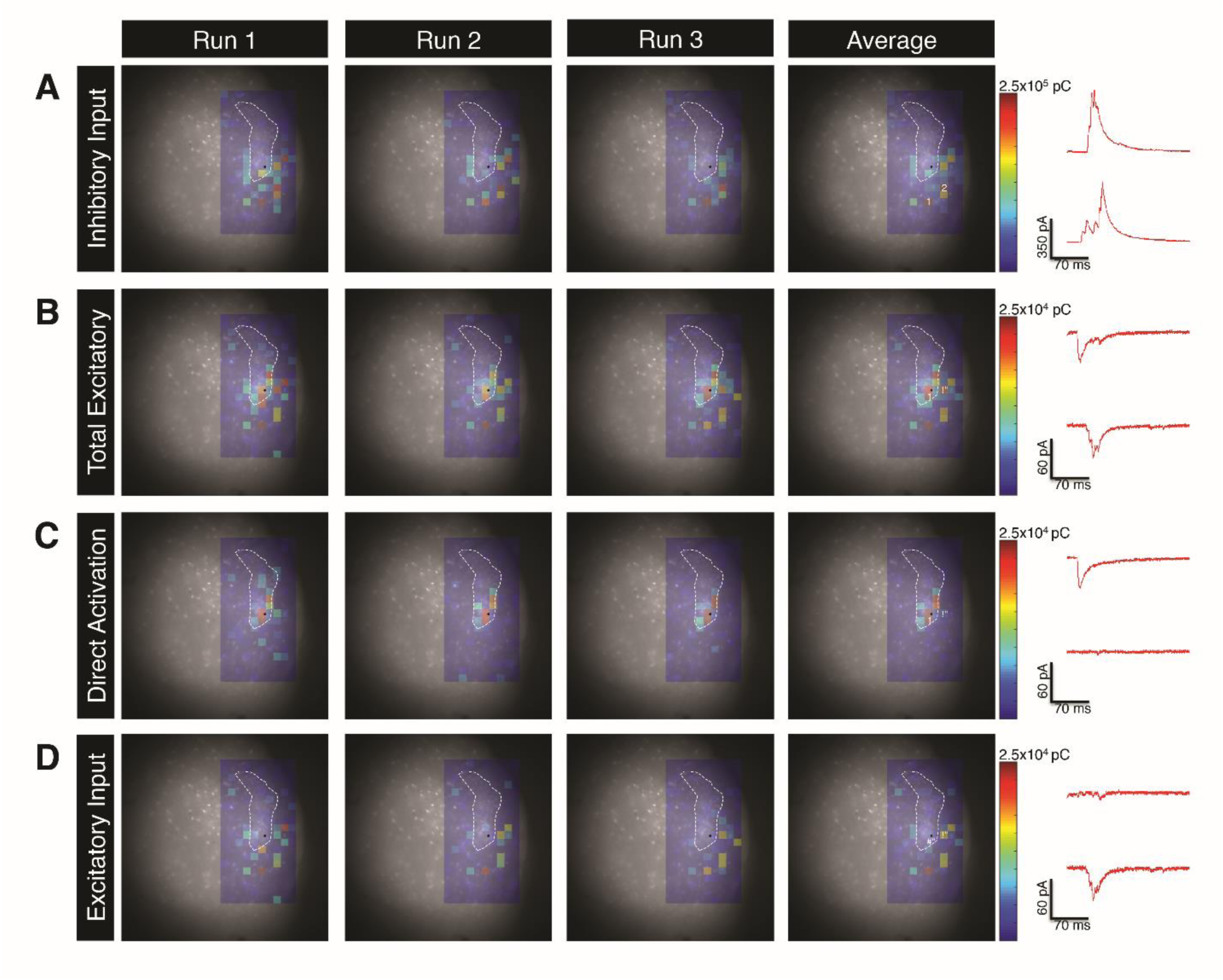

**Supplemental Figure 3:**
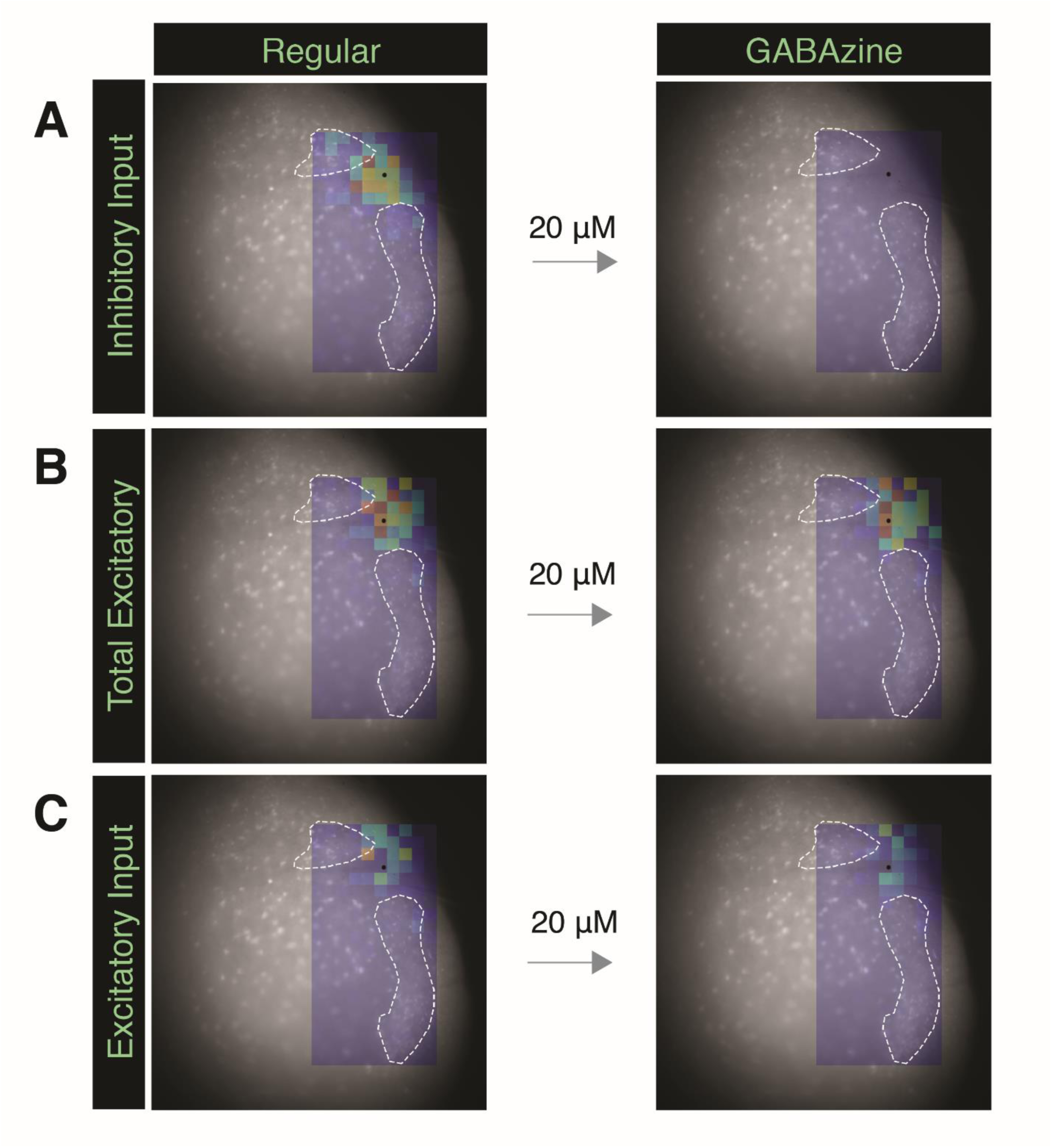
Examples of inhibitory and excitatory responses. A) Inhibitory responses for three separate runs. The average maps (as shown in the final panel) were analyzed and used for quantification. Example traces from two stimulation sites (1 = top trace, 2 = bottom trace) are shown to the right. B) Total excitatory responses, including both direct activation and synaptic input to the recorded cell. C) Excitatory responses in a low-calcium ACSF which blocks synaptic inputs, giving rise to only direct activation of the recorded cell. Note that the response in the bottom trace from B is no longer present, indicating that it was a synaptic input. D) Maps of the excitatory synaptic inputs were generated by subtracting the direct input maps from the total excitatory maps.

**Supplemental Figure 4:**
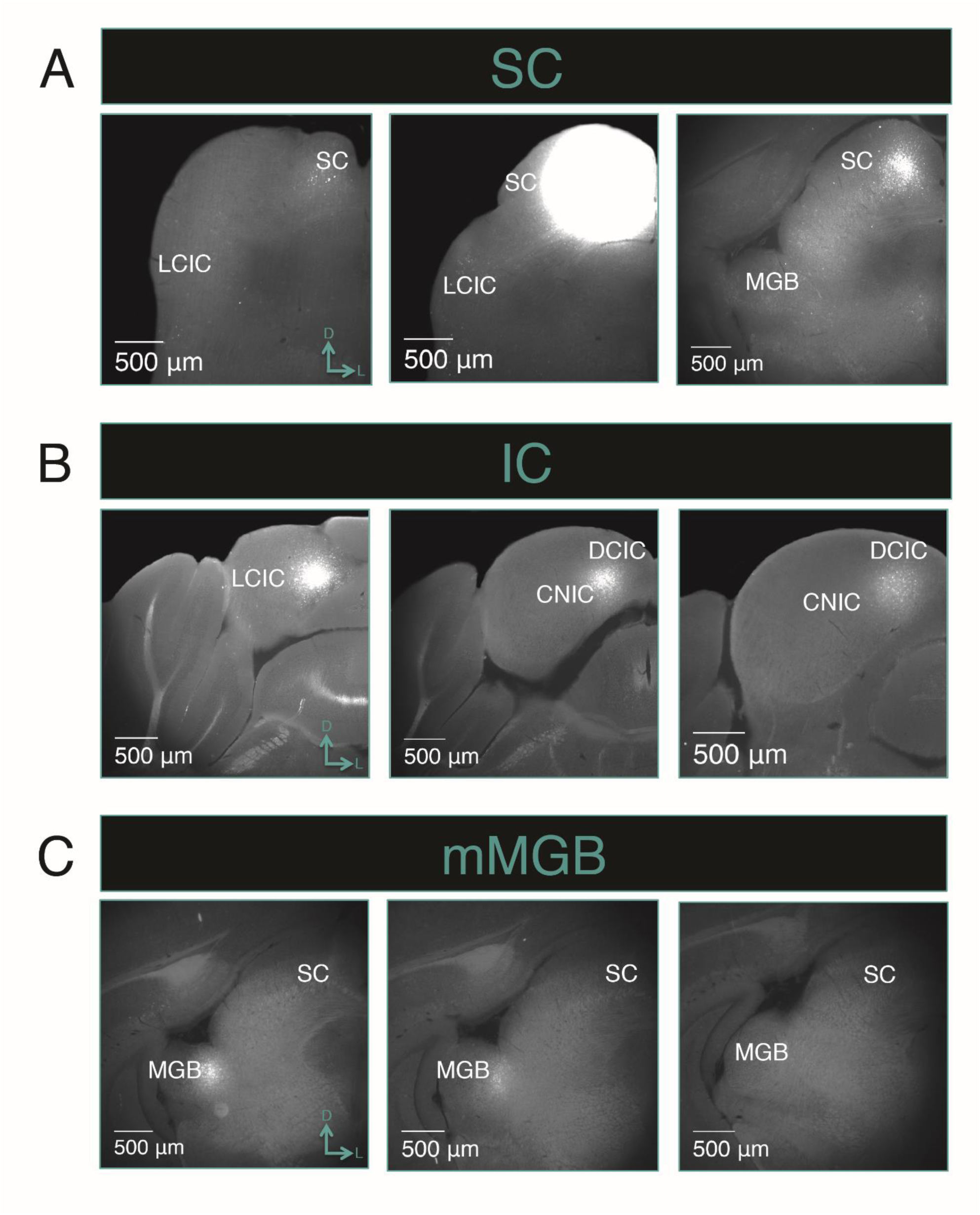
Injection sites. A) Injection site into the SC shown at multiple rostro-caudal levels (left panel: caudal, middle panel: mid-rostro-caudal, right panel: rostral). B) Injection site into the IC shown at multiple rostro-caudal levels. C) Injection site into the mMGB shown at multiple rostro-caudal levels.

## REFERENCES

Aitkin LM (1973) Medial geniculate body of the cat: responses to tonal stimuli of neurons in medial division. Journal of neurophysiology 36:275–283.

Aitkin LM, Kenyon CE, Philpott P (1981) The representation of the auditory and somatosensory systems in the external nucleus of the cat inferior colliculus. Journal of Comparative Neurology 196:25–40.

Aitkin LM, Dickhaus H, Schult W, Zimmermann M (1978) External Nucleus of Inferior Colliculus: Auditory and Spinal Somatosensory Merents and Their Interactions.

Andersen RA, Snyder RL, Merzenich MM (1980) The topographic organization of corticocollicular projections from physiologically identified loci in the AI, AII, and anterior auditory cortical fields of the cat. Journal of Comparative Neurology 191:479–494.

Bordi F, LeDoux JE (1994) Response properties of single units in areas of rat auditory thalamus that project to the amygdala. Experimental Brain Research 98:261–274.

Calford MB, Aitkin LM (1983) Ascending projections to the medial geniculate body of the cat: evidence for multiple, parallel auditory pathways through thalamus. Journal of Neuroscience 3:2365–2380.

Casseday JH, Fremouw T, Covey E (2002) The inferior colliculus: a hub for the central auditory system. In: Integrative functions in the mammalian auditory pathway, pp 238–318: Springer.

Chandrasekaran L, Xiao Y, Sivaramakrishnan S (2013) Functional architecture of the inferior colliculus revealed with voltage-sensitive dyes. Frontiers in neural circuits 7.

Chernock ML, Larue DT, Winer JA (2004) A periodic network of neurochemical modules in the inferior colliculus. Hearing Research 188:12–20.

Coleman JR, Clerici WJ (1987) Sources of projections to subdivisions of the inferior colliculus in the rat. Journal of Comparative Neurology 262:215–226.

Davis M, Gendelman DS, Tischler MD, Gendelman PM (1982) A primary acoustic startle circuit: lesion and stimulation studies. Journal of Neuroscience 2:791– 805.

Dillingham CH, Gay SM, Behrooz R, Gabriele ML (2017) Modular-extramodular organization in developing multisensory shell regions of the mouse inferior colliculus. Journal of Comparative Neurology.

Friauf E, Kandler K (1990) Auditory projections to the inferior colliculus of the rat are present by birth. Neuroscience letters 120:58–61.

Gabriele ML, Brunso-Bechtold JK, Henkel CK (2000a) Plasticity in the development of afferent patterns in the inferior colliculus of the rat after unilateral cochlear ablation. Journal of Neuroscience 20:6939–6949.

Gabriele ML, Brunso-Bechtold JK, Henkel CK (2000b) Development of afferent patterns in the inferior colliculus of the rat: projection from the dorsal nucleus of the lateral lemniscus. Journal of Comparative Neurology 416:368– 382.

Gao E, Suga N (2000) Experience-dependent plasticity in the auditory cortex and the inferior colliculus of bats: role of the corticofugal system. Proceedings of the National Academy of Sciences 97:8081–8086.

Gerfen CR (1984) The neostriatal mosaic: compartmentalization of corticostriatal input and striatonigral output systems. Nature 311:461–464.

Gerfen CR (1985) The neostriatal mosaic. I. Compartmental organization of projections from the striatum to the substantia nigra in the rat. Journal of Comparative Neurology 236:454–476.

Gerfen CR (1992) The neostriatal mosaic: multiple levels of compartmental organization. In: Advances in Neuroscience and Schizophrenia, pp 43–59: Springer.

Goldowitz D (2010) Allen Reference Atlas. A Digital Color Brain Atlas of the C57BL/6J Male Mouse-by HW Dong. Genes, Brain and Behavior 9:128–128.

González-Hernández T, Mantolán-Sarmiento B, González-González B, Pérez-González H (1996) Sources of GABAergic input to the inferior colliculus of the rat. Journal of Comparative Neurology 372:309–326.

Graybiel AM, Ragsdale CW (1978) Histochemically distinct compartments in the striatum of human, monkeys, and cat demonstrated by acetylthiocholinesterase staining. Proceedings of the National Academy of Sciences 75:5723–5726.

Huffman RF, Henson Jr OW (1990) The descending auditory pathway and acousticomotor systems: connections with the inferior colliculus. Brain Research Reviews 15:295–323.

Ikrar T, Olivas ND, Shi Y, Xu X (2011) Mapping inhibitory neuronal circuits by laser scanning photostimulation. Journal of visualized experiments: JoVE.

Jain R, Shore S (2006) External inferior colliculus integrates trigeminal and acoustic information: unit responses to trigeminal nucleus and acoustic stimulation in the guinea pig. Neuroscience letters 395:71–75.

Kincaid AE, Wilson CJ (1996) Corticostriatal innervation of the patch and matrix in the rat neostriatum. The Journal of comparative neurology 374:578–592.

Koch M, Schnitzler H-U (1997) The acoustic startle response in rats—circuits mediating evocation, inhibition and potentiation. Behavioural brain research 89:35–49.

LeDoux JE, Sakaguchi A, Reis DJ (1984) Subcortical efferent projections of the medial geniculate nucleus mediate emotional responses conditioned to acoustic stimuli. Journal of Neuroscience 4:683–698.

LeDoux JE, Ruggiero DA, Reis DJ (1985) Projections to the subcortical forebrain from anatomically defined regions of the medial geniculate body in the rat. Journal of Comparative Neurology 242:182–213.

Ledoux JE, Ruggiero DA, Forest R, Stornetta R, Reis DJ (1987) Topographic organization of convergent projections to the thalamus from the inferior colliculus and spinal cord in the rat. Journal of Comparative Neurology 264:123–146.

Lesicko AM, Hristova TS, Maigler KC, Llano DA (2016) Connectional modularity of top-down and bottom-up multimodal inputs to the lateral cortex of the mouse inferior colliculus. Journal of Neuroscience 36:11037–11050.

Llano DA, Sherman SM (2009) Differences in intrinsic properties and local network connectivity of identified layer 5 and layer 6 adult mouse auditory corticothalamic neurons support a dual corticothalamic projection hypothesis. Cerebral cortex 19:2810–2826.

Motts SD, Schofield BR (2009) Sources of cholinergic input to the inferior colliculus. Neuroscience 160:103–114.

Ono M, Yanagawa Y, Koyano K (2005) GABAergic neurons in inferior colliculus of the GAD67-GFP knock-in mouse: electrophysiological and morphological properties. Neuroscience research 51:475–492.

Petersen CC (2007) The functional organization of the barrel cortex. Neuron 56:339– 355.

Rouiller E, Rodrigues-Dagaeff C, Simm G, De Ribaupierre Y, Villa A, De Ribaupierre F (1989) Functional organization of the medial division of the medial geniculate body of the cat: tonotopic organization, spatial distribution of response properties and cortical connections. Hearing research 39:127–142.

Schofield BR (2010) Projections from auditory cortex to midbrain cholinergic neurons that project to the inferior colliculus. Neuroscience 166:231–240.

Schwarz C, Thier P (1995) Modular organization of the pontine nuclei: dendritic fields of identified pontine projection neurons in the rat respect the borders of cortical afferent fields. Journal of Neuroscience 15:3475–3489.

Shepherd GM (2012) Circuit mapping by ultraviolet uncaging of glutamate. Cold Spring Harb Protoc 2012:998–1004.

Shepherd GM, Pologruto TA, Svoboda K (2003) Circuit analysis of experiencedependent plasticity in the developing rat barrel cortex. Neuron 38:277–289.

Sivaramakrishnan S, Oliver DL (2006) Neuronal responses to lemniscal stimulation in laminar brain slices of the inferior colliculus. Journal of the Association for Research in Otolaryngology 7:1–14.

Stebbings KA, Lesicko AM, Llano DA (2014) The auditory corticocollicular system: molecular and circuit-level considerations. Hear Res 314:51–59.

Sturm J, Nguyen T, Kandler K (2014) Development of intrinsic connectivity in the central nucleus of the mouse inferior colliculus. The Journal of neuroscience: the official journal of the Society for Neuroscience 34:15032–15046.

Swerdlow N, Geyer M, Braff D (2001) Neural circuit regulation of prepulse inhibition of startle in the rat: current knowledge and future challenges. Psychopharmacology 156:194–215.

Syka J, Straschill M (1970) Activation of superior colliculus neurons and motor responses after electrical stimulation of the inferior colliculus. Experimental neurology 28:384–392.

Tamamaki N, Yanagawa Y, Tomioka R, Miyazaki JI, Obata K, Kaneko T (2003) Green fluorescent protein expression and colocalization with calretinin, parvalbumin, and somatostatin in the GAD67-GFP knock-in mouse. Journal of Comparative Neurology 467:60–79.

Winer JA (1992) The functional architecture of the medial geniculate body and the primary auditory cortex. In: The mammalian auditory pathway: Neuroanatomy, pp 222–409: Springer.

Winer JA, Larue DT, Diehl JJ, Hefti BJ (1998) Auditory cortical projections to the cat inferior colliculus. Journal of Comparative Neurology 400:147–174.

Wu C, Stefanescu RA, Martel DT, Shore SE (2014) Listening to another sense: somatosensory integration in the auditory system. Cell and tissue research: 1– 18.

